# The PIWI/piRNA response is relaxed in a rodent that lacks mobilizing transposable elements

**DOI:** 10.1101/587030

**Authors:** Michael W. Vandewege, Roy N. Patt, Dana K. Merriman, David A. Ray, Federico G. Hoffmann

## Abstract

Transposable elements (TEs) are genomic parasites that can propagate by inserting copies of themselves into host genomes. Mammalian genomes are typically dominated by LINE retrotransposons and their associated SINEs, and their mobilization in the germline is a challenge to genome integrity. There are genomic defenses against TE proliferation and the PIWI/piRNA defense is among the most well understood. However, the PIWI/piRNA system has been investigated largely in animals with abundant and actively mobilizing TEs and it is unclear how the PIWI/piRNA system functions in the absence of mobilizing TEs. The 13-lined ground squirrel provides an excellent opportunity to examine PIWI/piRNA and TE dynamics within the context of minimal, and possibly nonexistent, TE accumulation. We sequenced RNA and small RNAs pools from the testes of juvenile and adult squirrels and compared results to TE and PIWI/piRNA dynamics in the European rabbit and house mouse. Interestingly in squirrels, despite a lack of young insertions, TEs were still actively transcribed at higher levels compared to mouse and rabbit. All three PIWI proteins were either not expressed, or only minimally expressed, prior to P8 in squirrel testis, but there was little TE expression change with the onset of PIWI expression. We found PIWIs largely did not reduce TE transcription, and the ping-pong cycle was significantly reduced among squirrel LINEs and SINEs compared to the mouse and rabbit. We speculate that, although the PIWI/piRNA system is adaptable to novel TE threats, transcripts from TEs that are no longer threatening receive less attention from PIWI proteins.

## Introduction

Transposable elements (TEs) are genomic parasites that propagate by inserting copies of themselves into the genomes of their hosts. They account for up to 70% of mammalian genome content (de Koning et al. 2011). Because of their ability to mobilize, TEs are powerful mutagens, as novel TE insertions can disrupt exons, regulatory elements, and splice junctions, and facilitate non-homologous recombination. As a result, TE insertions have been linked to genomic deletions, duplications, inversions, translocations, and chromosome breaks in a variety of genomes (Cheng et al. 2005; Franke et al. 2017; Platt et al. 2018). While some TE insertions have proven adaptive, TEs are generally considered a serious challenge to genome integrity.

Eukaryotic genomes have evolved mechanisms to restrict TE mobilization, especially in the germline. PIWI proteins and PIWI-interacting RNAs (piRNAs) have emerged as key components in protecting the genome against the proliferation of TEs, and probably evolved in response to the challenge presented by them (Aravin, Hannon, et al. 2007; Brennecke et al. 2007; Aravin et al. 2008; Malone and Hannon 2009; Siomi et al. 2011). piRNAs and PIWI proteins assemble into RNA-induced silencing complexes, which go on to neutralize TE-like targets by transcript cleavage or chromatin methylation (Aravin, Sachidanandam, et al. 2007; Carmell et al. 2007; Houwing et al. 2007; Molaro et al. 2014).

Of the characterized PIWI/piRNA models, vertebrates, fruit fly, and *C. elegans,* each possess independently derived sets of PIWI paralogs which in turn associate with different sets of piRNAs (Reddien et al. 2005; Kerner et al. 2011; Lewis et al. 2016). In mammals, mouse (*Mus musculus*) is the most studied system (reviewed in Ernst et al. 2017). Mouse piRNAs have been categorized into two major sets, pre-pachytene and pachytene, according to when their expression begins (Aravin et al. 2006; Girard et al. 2006). Pre-pachytene piRNAs are first expressed in the early stages of spermatogenesis, are enriched for TE-like sequences, and preferentially associate with the PIWIL2 and PIWIL4 paralogs. These piRNAs are thought to be largely responsible for silencing TEs. By contrast, pachytene piRNAs begin their expression when spermatocytes reach the pachytene stage, are largely derived from intergenic regions, and preferentially associate with PIWIL1. Even though pachytene piRNAs are the most abundant piRNA in adult mouse testes, their biological role is still poorly understood and current evidence suggests they play a role similar to microRNAs, silencing messenger RNAs in mouse spermiogenesis (Li et al. 2013; Gou et al. 2014; Wu et al. 2020).

The mouse genome encodes three PIWI paralogs that are differentially expressed during development and associate preferentially with piRNAs of different sizes. PIWIL2, also known as MILI, preferentially binds piRNAs that are 26-27 nucleotides (nts) long, begins expression at embryonic day 12.5 (E12.5) in developing testes, and is linked to the post-transcriptional silencing of TEs. PIWIL4, or MIWI2, preferentially binds to piRNAs that are ~28 nts long, is expressed between E15.5 and post-natal day 3 (P3), and this period of time is linked to the *de novo* establishment of methylation marks in primordial germ cells (Aravin et al. 2008). Finally, PIWIL1, or MIWI, begins expression at P14, when the pachytene stage of meiosis begins and continues through adulthood. This paralog preferentially binds to piRNAs ~30 nts long and is mostly linked to spermiogenesis (Lau et al. 2006). Both PIWIL2 and PIWIL4 participate in a feed-forward loop known as the ping-pong cycle that serves to reduce the abundance of TE transcripts through piRNA-guided cleavage of TE-derived mRNA and methylation of TE loci (Aravin, Sachidanandam, et al. 2007; Aravin et al. 2008; Kuramochi-Miyagawa et al. 2008; De Fazio et al. 2011; Manakov et al. 2015). A complete understanding of the functional roles of piRNAs is still lacking, but we know that defects in the PIWI paralogs lead to elevated TE activity and problems in the male germline (Carmell et al. 2007; Houwing et al. 2007; O’Donnell and Boeke 2007).

TE expression and accumulation rates vary among different mammals, both in terms of the type of TEs that are active and in the level of challenge presented by them (Pasquesi et al. 2020). However, piRNAs have only been described in a handful of vertebrates and a comparative framework is generally lacking (Lau et al. 2006; Liu et al. 2012; Chirn et al. 2015; Toombs et al. 2016; Vandewege et al. 2016; Sun et al. 2017). Further, whether the developmental changes in PIWI and piRNA expression are conserved, or how PIWIs/piRNAs repertoires respond to changing TE landscapes has not been addressed. In this regard, there is evidence that the PIWI/piRNA system is quickly adaptable to novel TE threats (Mourier 2011; Gainetdinov et al. 2017; Sun et al. 2017; Zhang et al. 2020), but it is not clear what happens when established TEs are no longer mobilizing or threatening the genome.

Comparisons among the 13-lined ground squirrel (*Ictidomys tridecemlineatus*), rabbit (*Oryctolagus cuniculus*), and mouse (*Mus musculus*) offer a natural experiment to test these questions. Genomic surveys indicate that the last LINE and SINE insertions in the 13-lined ground squirrel genome occurred between four and five million years ago and no retrotransposition-competent LINE loci have been found in its genome so far (Platt and Ray 2012). Thus, the squirrel offers a unique opportunity to study piRNA biogenesis, piRNA diversity, and the ping-pong cycle in a system where LINE and SINE mobilization appears to have ceased. By contrast, mouse and rabbit have typical mammalian genomic TE landscapes dominated by currently active retrotransposons, LINE-1 (L1), and associated SINEs, but have different patterns of TE expression and accumulation from one another. In the current study, we took advantage of the genomic resources available for these three species to 1- explore the relationship between TE abundance at the genome and transcript level in each species, 2- characterize developmental changes in the expression of PIWI paralogs and piRNAs, 3- measure the intensity of the ping-pong cycle and, 4- quantify TE expression changes in response to changing PIWI expression. We found that TE expression was much higher in the squirrel, despite the observed absence of recent TE insertions, but that the piRNA/PIWI response was reduced. Our results suggest that TEs that lack the capacity to generate *de novo* insertions elicit weak responses from the piRNA/PIWI pathway.

## Results

### TE accumulation

We first characterized patterns of TE accumulation, diversity, and abundance in the mouse, rabbit, and squirrel genomes and found clear differences among them. We assessed the abundance and diversity of TEs present in each genome by measuring overall insertion numbers, young insertion numbers (<5% diverged from consensus), and median family distances. As in most mammals, LINEs, SINEs, and LTR retrotransposons account for the vast majority of TE insertions in these three species. Close to half of the TE-derived portion of the genome corresponds to LINEs, with a low number of insertions from DNA transposons in all three genomes (Fig. 1A). In contrast, there were differences in the relative contribution of SINEs and LTRs among them: SINEs accounted for close to half of the TE-derived portion of the genome in the rabbit, and LTR retrotransposons contributed a larger fraction in squirrel and mouse relative to the rabbit. These three species also differed in the historical patterns of TE accumulation. To reconstruct TE deposition history, we classified insertions by subfamilies and estimated the Kimura 2-parameter distance between consensus sequences of each TE family and individual insertions. According to the master gene model, TE insertions are driven by one master mobilizing element (Deininger et al. 1992; Cordaux et al. 2004), and younger insertions would more closely resemble the corresponding master mobilizing element than older insertions. Thus, distances to the corresponding consensus can be used to estimate the relative age of an insertion. In the case of the squirrel, these analyses confirmed a low number of young TE insertions, as previously reported, which would indicate a lack of mobilizing elements (Fig. 1B; Platt and Ray 2012).

**Figure 1.**
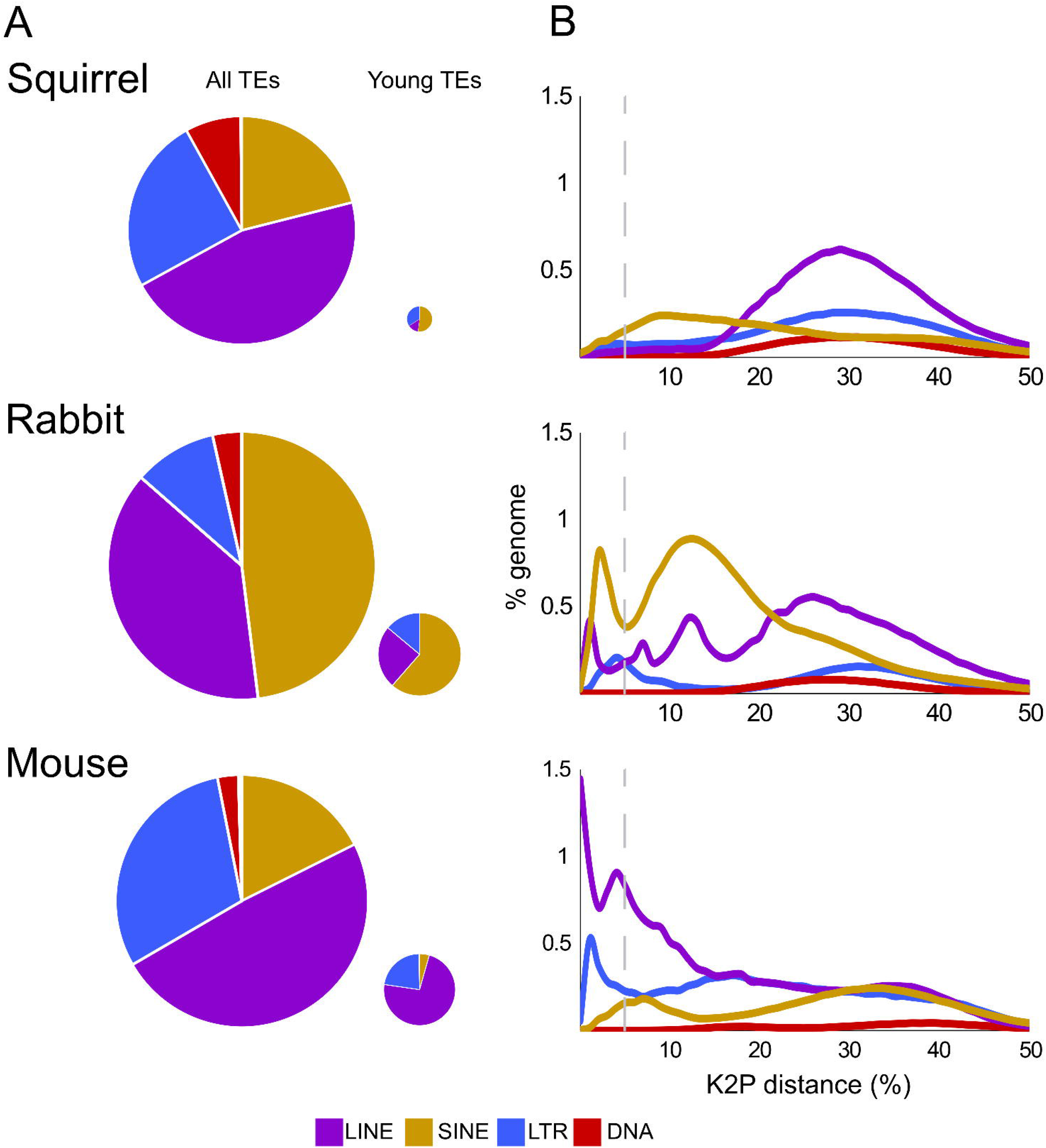
A) Relative contribution of TEs to the corresponding genome. The small pie charts document the contribution of the more recent TE insertions, with the size of the pie chart proportional to the relative contribution of recent insertions to each genome. B) Historical patterns of accumulation of the major TE categories inferred by calculating the Kimura 2-parameter distance between individual insertions and the corresponding consensus. Relatively young (<5% diverged) insertions to the left of the vertical line with lower genetic distances were deposited more recently. The insertions to the left of the dashed vertical lines were considered for the small pie charts.

### PIWI expression and piRNAs

We then assessed changes in the expression of PIWI genes and associated piRNAs within developing testes in squirrel and rabbit and compared them to publicly available data from mouse. We conducted simultaneous RNA-Seq and small RNA-Seq experiments on a neonate (P0 – 0-days post-birth), post-natal juveniles (P2, P8, P10, P13), and one adult in the case of the squirrel, whereas in rabbits, we sampled testes from a P21 juvenile and one adult (P > 90). As expected, we observed changes in the expression of the PIWI paralogs and corresponding piRNA repertoires associated with developmental stage. In squirrel, we did not detect the expression of any PIWI paralog in the P0 and P2 samples, and the small RNAs libraries exhibited no clear hallmarks of PIWI processing (Fig 2A; Supplemental Fig 1A). After P8, all three paralogs were expressed. PIWIL4 was the most expressed, and piRNA biogenesis became apparent (Fig 2A; Supplemental Fig1A). The expression of squirrel PIWIL4 was unique because this gene was not expressed in our rabbit or mouse samples, and it was expressed in adult testis, which has not been previously recorded (Fig 2A). In all three species, piRNAs became more abundant and diverse with the onset of PIWIL1 expression in adult testis (Fig 2A, Supplemental Table 1). As PIWIL1 became the most abundantly expressed paralog, the size distribution of piRNAs shifted in mouse and rabbit from a peak length of 27-28 nts in juveniles to a peak of 29-30 nts in adults (Fig. 2A). In contrast, the piRNA length distribution in adult squirrels remained centered at 28 nts, likely due to the continued expression of PIWIL4 which tends to produce shorter piRNAs than PIWIL1.

**Figure 2.**
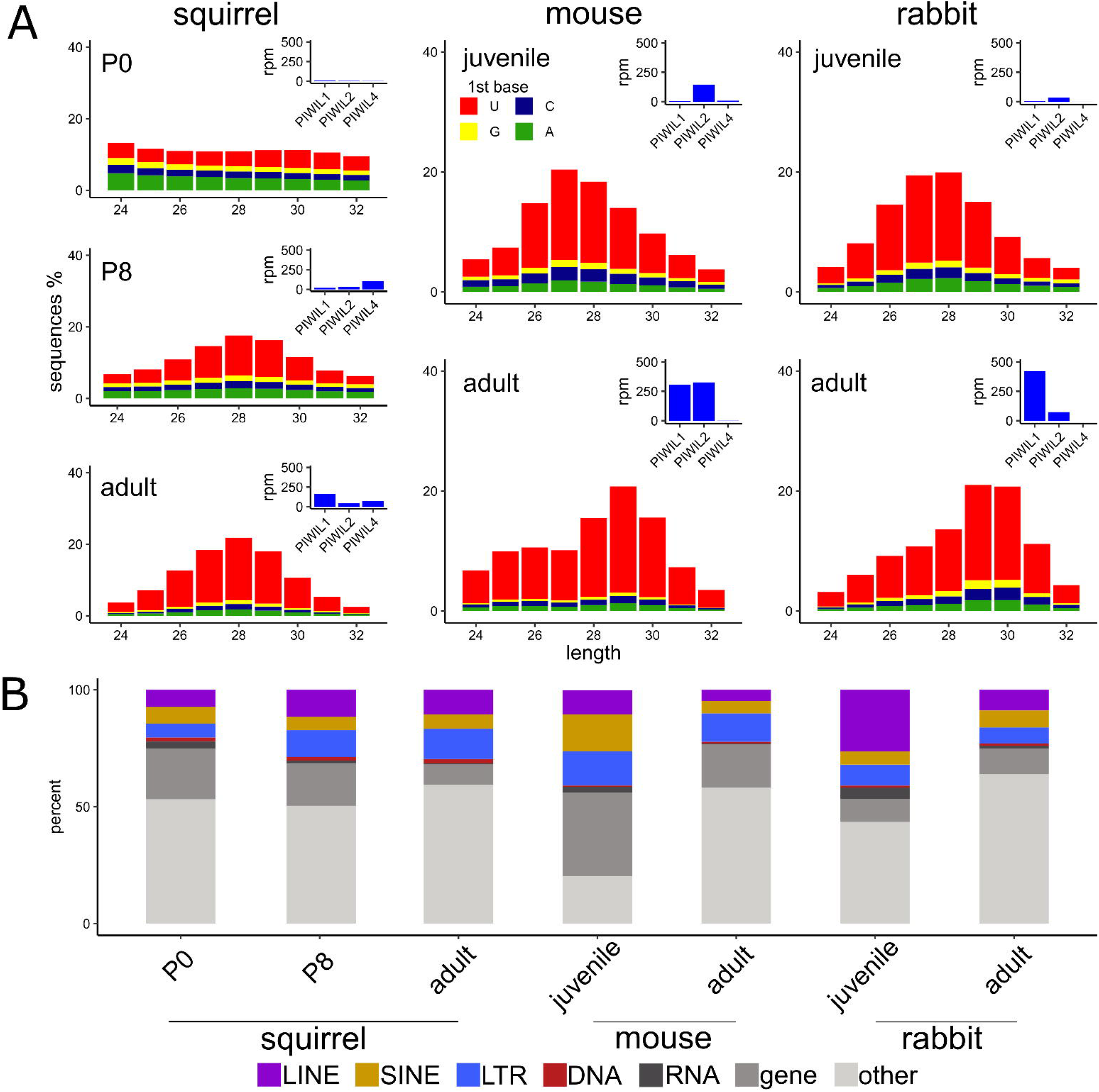
A) Distribution length among piRNAs between 24 and 32 nt. Colors in the stacked bar plot represent the frequency of the first nucleotide. Corresponding expression of the PIWI proteins is measured as reads per million in each sample. B) The proportion of piRNAs that mapped to discrete regions in the genome.

TEs are known to be a more common source of prepachytene piRNAs in juvenile testis, and pachytene piRNAs in adult testis are mostly derived from intergenic space. This was observed in both the rabbit and mouse, but was not as prevalent in squirrel (Fig 2B). In squirrel, TEs became a more common source of the small RNAs once PIWIs were expressed, but TE-derived piRNAs were almost equally frequent in juvenile and adult testis (Fig 2B).

### TE expression

We mapped RNA-Seq reads to corresponding genomes and measured TE family expression in units of reads per million (RPM). This metric can be expressed as a percent of the total RNA pool, which allows for comparisons across species. We found that squirrel TEs made up more than 15% of the transcriptome, almost doubling the relative amount of TE-derived RNA obtained from the corresponding mouse and rabbit samples (Fig 3A). When we examined changes in TE expression over time, we noticed some general changes between the juvenile and adult testes of mouse and rabbit, but found that changes in TE expression during the development of squirrel testis were less dramatic. For example, LINE expression was lower in the testes of juvenile mouse and rabbit, compared to adult, but was roughly equivalent in all stages of squirrel samples examined (Supplemental Fig 2A). Interestingly, the percent of the transcriptome made up of young TEs was similar across all species and developmental stages, with the exceptions of the juvenile rabbit and adult squirrel (Fig 3A). These exceptions were largely caused by the high expression of a few LTR and SINE families (Supplemental Fig 2B). The mouse genome contains numerous young LINE families that are actively mobilizing (Goodier et al. 2001; Sookdeo et al. 2013) and most of the young TEs from these families were expressed. By contrast, most young insertions in the rabbit and squirrel were generated by just two families that accounted for most of the RNA derived from young LINEs (Supplemental Fig 2B).

**Figure 3.**
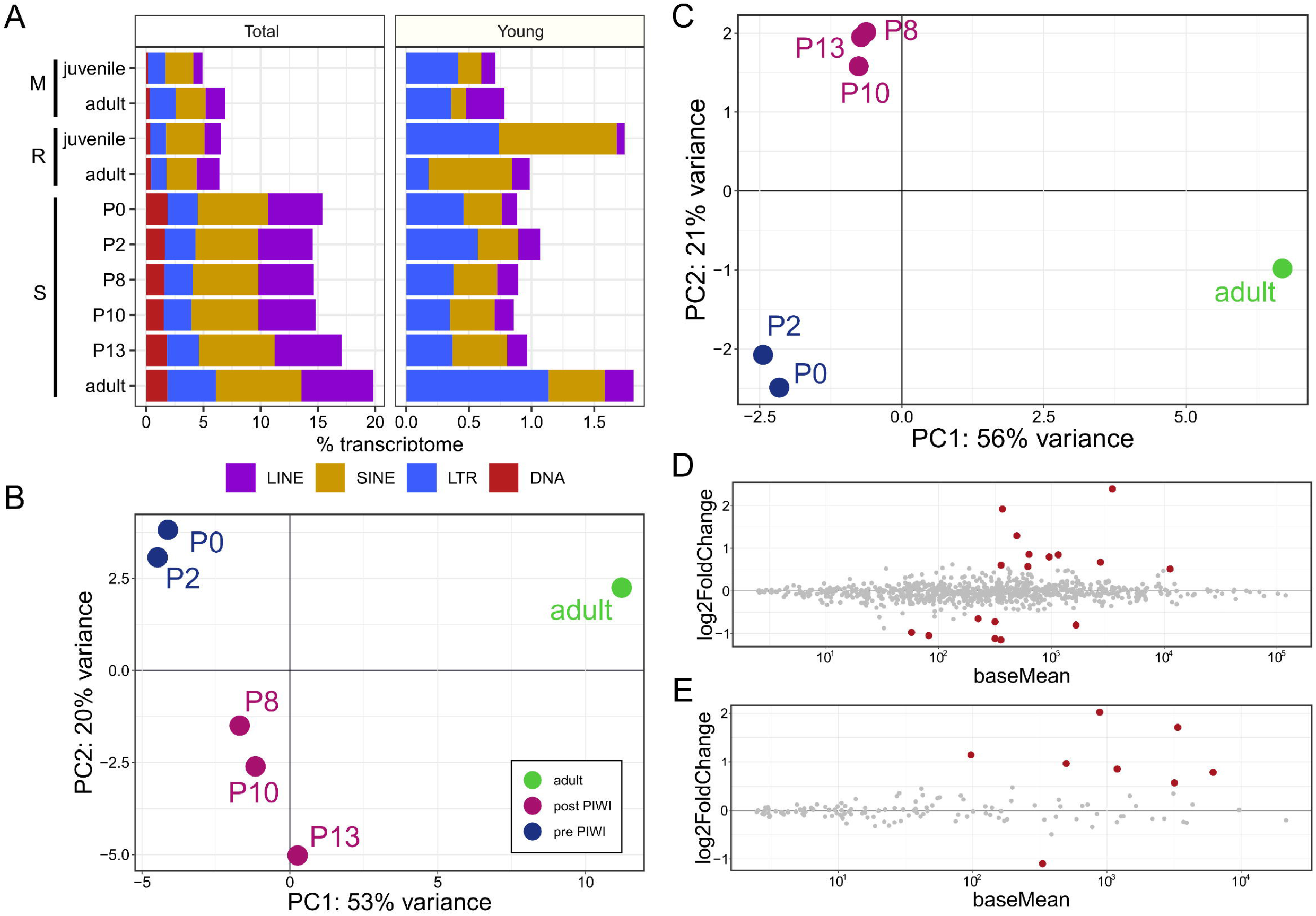
A) TE expression among all samples measured as the percent transcriptome among all TE insertions and insertions < 5% diverged from the consensus (young). S, m, and r reflect samples from the squirrel, mouse, and rabbit, respectively. PCA of squirrel testis derived from the expression of all TE insertions (B) and young insertions (C). Differentially expressed squirrel TEs between the pre-PIWI expressed samples (P0, P2) and post-PIWI expressed juvenile samples (P8, P10, P13) calculated from all TE insertions (D) and young insertions (E). Red circles reflect differentially expressed TEs with an adjusted p-value < 0.1. Positive log2fold change values reflect higher expression in pre-PIWI samples.

Since multiple life stages from the squirrel were sampled, we examined variation in TE expression among these developmental stages. We first conducted a principal component analysis (PCA) from the expression of all genes and TE insertions. We found that the first two principal components accounted for 90% of the variation and samples clustered into three groups that reflect the observed changes in PIWIL and piRNA expression. The P0 and P2 samples that lacked expressed PIWIs and piRNAs clustered in one group, the juvenile P8, P10 and P13 samples with expressed PIWIs and piRNAs clustered in another, and the adult sample was isolated (Supplemental Fig3A). When we excluded gene expression data, we found that samples grouped in a similar manner whether we considered expression from all TEs (Fig 2B), or only young insertions (Fig 2C), although samples were closer together in principle component space compared to the whole expression profile. Because the grouping of these clusters corresponded well with the observed changes in PIWI expression, we used these life stage replicates to test whether TE expression changed with the onset of PIWI paralog expression. We performed a differential expression analysis between the pre-PIWI and post-PIWI juvenile samples (P0 and P2 vs. P8, P10, and P13) to determine if TEs were less expressed in post-PIWI samples -- as the ping-pong model would predict. However, TE expression was similar among samples, and only 17 out of 794 families were differentially expressed. There was an almost equal mix of up- and down-regulated TE families: 10 families were more expressed in pre-PIWI samples whereas seven were more expressed in the post-PIWI samples (Fig 2D). Further, log2 fold expression was less than two in all but one family. Most of these 17 families had median K2P distances larger than 0.1 and were mostly LTRs (Supplemental Fig 3B). We repeated this analysis for just the young insertions and found that eight out of 151 families with young insertions were differentially expressed, and that the log2 fold change was less than two in all cases (Fig 2E). All differentially expressed families were LTRs and six were relatively young, with median K2P distances < 0.1 (Supplemental Fig 3C). Seven out of the eight families were upregulated in pre-PIWI samples, suggesting PIWIs had a stronger effect on these young insertions compared to all insertions, as the ping-pong cycle would predict. However, very few families were differentially expressed, fold changes were relatively low, and only LTR insertions exhibited decreased expression with the onset of PIWI expression. These results suggest TE transcription is not heavily regulated in squirrel testis.

### piRNAs and TE dynamics

Our next step was to compare how PIWI and piRNA targeting among TE families varied among the three species. We estimated the strength of the piRNA/PIWI response to TEs by calculating Z_10_ scores from mapped piRNAs in LINEs, SINEs and LTRs in all samples, for all insertions and for only young insertions. The Z_10_ score measures the frequency of complementary piRNAs that overlap by 10 nts and reflects the strength of the ping-pong cycle. A Z_10_ score > 1.96 reflects a non-random bias of complementary piRNAs overlapping by 10 bp and a p-value < 0.05. When we compared Z_10_ scores among squirrel samples, our results were consistent with measures of PIWI expression. piRNA overlaps were largely random or nonexistent among piRNAs mapping to TE families in the P0 and P2 squirrel testes, and the ping-pong signature only became noticeable with the onset of PIWI expression in P8. However, the majority of TE families still lacked a strong ping-pong signature (Supplemental Fig 4). To directly compare ping-pong cycle intensities across species, we averaged Z_10_ scores among samples within a species, but excluded the squirrel P0 and P2 samples because they lack PIWI proteins and piRNAs. An ANOVA revealed significant differences in Z_10_ score among species, so we conducted a post-hoc Tukey’s HSD test for pairwise comparisons and found that scores were significantly lower among squirrel LINEs than mouse and rabbit LINEs (p < 0.0001), Z_10_ scores were also significantly lower in squirrel SINEs compared to the mouse (p = 0.03), but not the rabbit (p = 0.45) (Fig 4A). We then examined Z_10_ scores among young insertions, but only compared young TE Z_10_ scores from families that had > 10 insertions < 5% diverged from consensus and non-zero Z_10_ scores. The same overall pattern was present, except Z_10_ scores in squirrel SINEs were lower than both mouse and rabbit (p < 0.05) (Fig 4B). The only TEs that had Z_10_ scores that reflected strong PIWI processing in the squirrel were the young LTRs (Fig 4B), consistent with our differential expression results (Supplemental Figure 3C).

**Figure 4.**
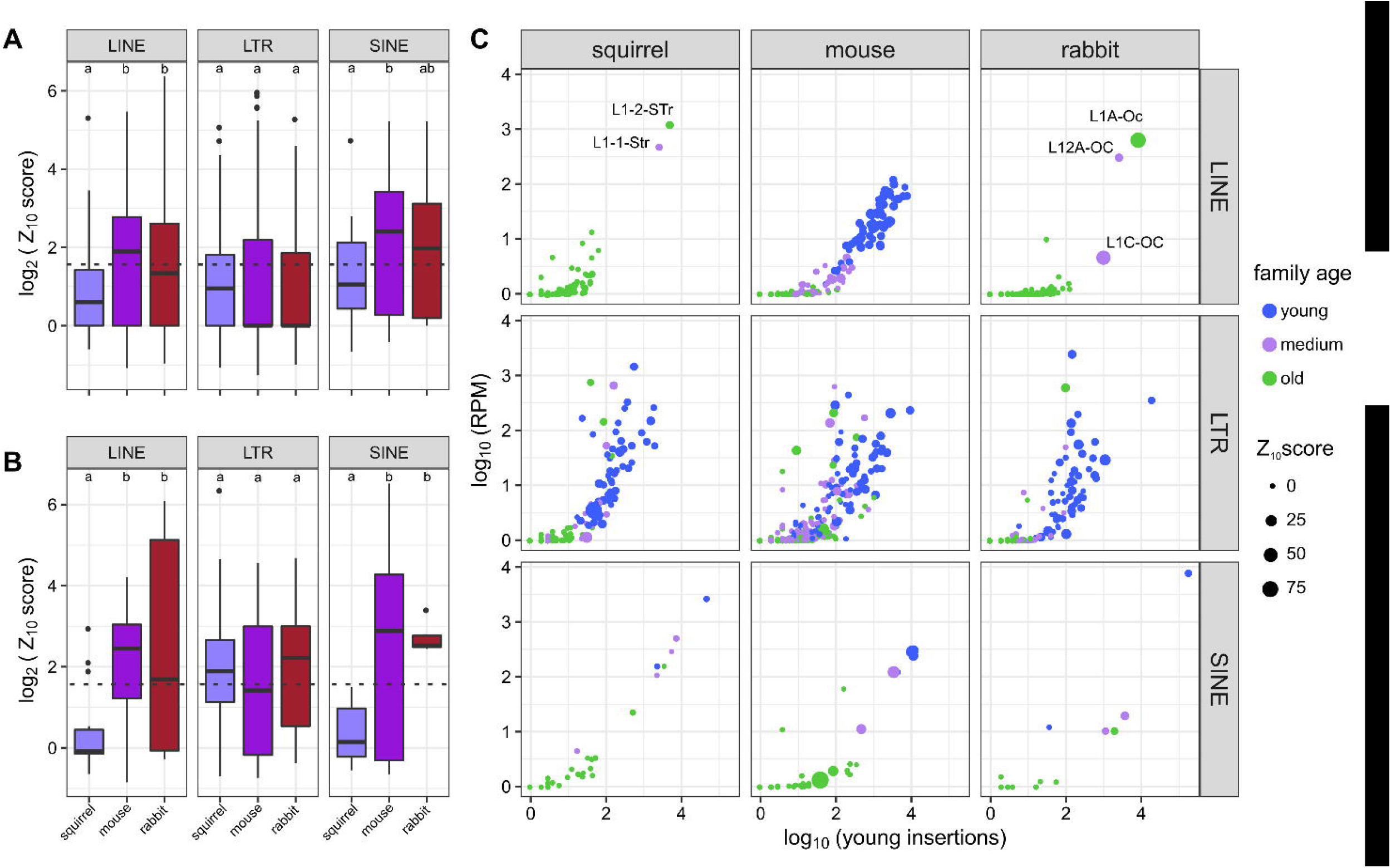
A) Z_10_ score distribution among families among species. The dotted line reflects the position of a Z_10_ score = 1.96, p = 0.05. Significant differences among species means (as determined by Tukey’s HSD test, p < 0.05) are indicated by different shared letters. B) Z_10_ score distributions calculated from young insertions. C) Number of young insertions plotted against the expression calculated from those insertions. Families are colored based on median K2P divergence, where young families are < 10%, medium are between 10% and 20% diverged, and old families are > 20%. Z_10_ scores calculated from young insertions are reflected by the size of the point.

We then compared average TE expression and Z_10_ scores to examine species-specific differences in TE and piRNA dynamics. First, we found that genome wide TE expression strongly correlated with the total number of insertions, and that the age of the TE family was less relevant. In fact, the most expressed families were older than 20% diverged from consensus, and almost every family had some expressed insertions (Supplemental Fig 5). Interestingly, Z_10_ scores did not strongly correlate with TE expression (Supplemental Fig 6), suggesting that PIWI proteins are somewhat selective and do not merely target the families with the highest expression. For example, young mouse LINE families (<10% median divergence) elicited a stronger response from PIWIs than did older families, regardless of expression. When we examined expression values and Z_10_ scores from young insertions, there was a strong positive correlation between insertion number and expression with a family age component, i.e. older families tended to have fewer young insertions that were less expressed (Figure 4C). The mouse LINEs held most closely to this pattern, *i.e.* young families with many young insertions were most expressed and exhibited more PIWI processing than insertions from medium-aged and old families. However, the two most active LINEs in squirrel and rabbit exhibited similar insertion number and expression values, but a dramatic difference in Z_10_ scores (Fig 4C). Regardless of whether we examined all or young insertions, PIWI proteins were not targeting LINEs and SINEs in the squirrel as heavily as in mouse or rabbit, but it was apparent that squirrel LTRs were still mobilizing and cleaved by PIWI proteins.

PIWI proteins can reduce the number of TE transcripts through direct cleavage and this targeting is reflected in the piRNA repertoire. piRNAs tend to derive from different TE sources in juvenile testis compared to adult testis, where LINEs are biased in juveniles and SINEs are biased in adults (Mourier 2011), suggesting a change in PIWI target preference. The ping-pong model predicts an inverse relationship between ping-pong cycle intensity and TE expression, *i.e.* if a TE is heavily cleaved by PIWIs, piRNAs should be abundant and expression should be lower; and vice versa. Therefore, we measured the log2 fold change in expression between juvenile (P13 of the squirrel) and adult testis, as well as the log2 fold change in Z_10_ scores in LINEs, and tested this prediction in the context of species-specific differences. We were particularly interested to find out how much PIWI proteins reduced the expression of the two most recent squirrel LINE families in juvenile testis. We examined only families with more than 200 young insertions, as this excludes the older LINEs in squirrel and rabbit that lack Z_10_ scores but still provides 68 data points from mouse to produce a distribution for comparison. The overall prediction was accurate, as most LINEs in juvenile testis exhibited higher Z_10_ scores and lower expression (Fig 5A). There was little expression or Z_10_ score change in the two squirrel LINEs, and we observed only a weak inverse relationship in one family when we examined these metrics from the youngest insertions (Fig 5B). These analyses indicate that, while these two L1s have several thousand young insertions and have Z_10_> 2.3 in all samples except P0 and P2, the strength of the ping-pong cycle in squirrel is not as intense as the cycle impacting active L1s in mouse or rabbit, and the PIWIs are not changing L1 expression.

**Figure 5.**
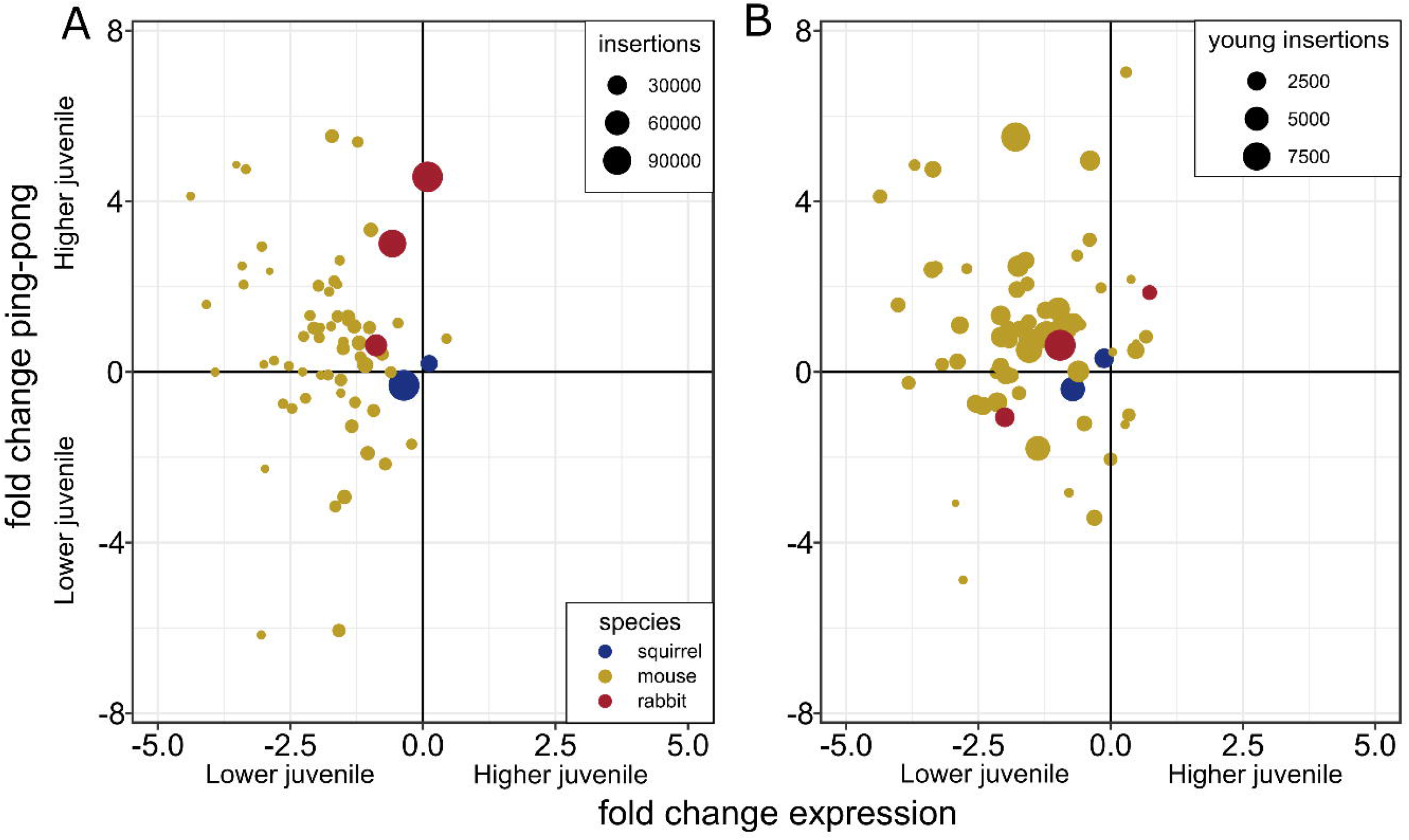
Changes in expression and the ping-pong cycle between a juvenile and adult for LINEs with more than 200 young insertions for A) all insertions for that family and B) just the young insertions. The juvenile squirrel is represented by the P13 sample.

### PIWI evolution

Finally, we tested if reduced PIWI targeting could be attributed to the PIWI proteins themselves, i.e. is there evidence of reduced function in the squirrel PIWI proteins due to deleterious mutations? To do so, we used selection tests that measure changes in d_N_/d_S_ (*ω*). We specifically used the RELAX module in HyPhy that tests whether the strength of natural selection (positive or purifying) has been relaxed or intensified along a specified test branch. We constructed trees from the PIWI sequences of Glires (Supplemental Figure 7) and used the squirrel branch as the test branch. This test was not significant for PIWIL2 (K = 1.28, p = 0.321, likelihood ratio = 0.99) or PIWIL4 (K = 0, p = 0.207, likelihood ratio = 1.59), but there was statistical support for the intensification of selection in PIWIL1 (K = 1.49, p = 0.03, likelihood ratio = 4.69). However, *ω* was less than 1 among reference branches in the alternative model, and a K = 1.49 pushed *ω* in the test branch closer to 0, indicating an intensification of purifying selection (Table 1). This suggests that PIWIs have not experienced relaxed selection but have maintained their functional constraint, such that the reduced targeting of squirrel LINEs and SINEs cannot be attributed to changes in PIWI function.

**Table 1.**
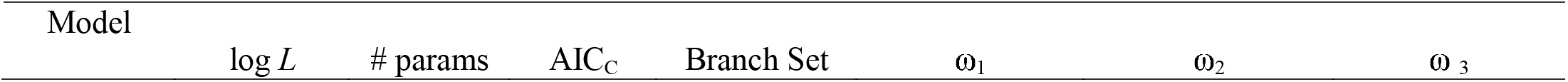
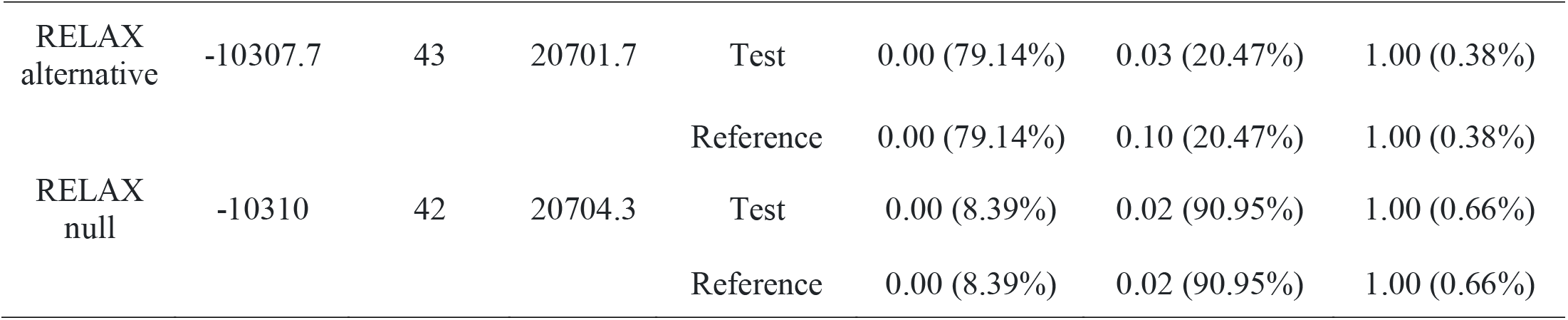
Results of RELAX tests for PIWIL1 on the squirrel branch.

| Model | log $L$ | # params | AIC<sub>C</sub> | Branch Set | $\omega_1$ | $\omega_2$ | $\omega_3$ |
| --- | --- | --- | --- | --- | --- | --- | --- |
| RELAX<br>alternative | -10307.7 | 43 | 20701.7 | Test | 0.00 (79.14%) | 0.03 (20.47%) | 1.00 (0.38%) |
|  |  |  |  | Reference | 0.00 (79.14%) | 0.10 (20.47%) | 1.00 (0.38%) |
| RELAX<br>null | -10310 | 42 | 20704.3 | Test | 0.00 (8.39%) | 0.02 (90.95%) | 1.00 (0.66%) |
|  |  |  |  | Reference | 0.00 (8.39%) | 0.02 (90.95%) | 1.00 (0.66%) |

## Discussion

TE landscapes are variable among vertebrates and, given that there are several interactions between TEs and host defenses, what exactly causes this variability is still an unanswered question in genomics. Here we attempted to quantify the contribution of PIWI proteins to the dearth of novel TE insertions in the squirrel genome by comparing TE accumulation, TE expression, PIWI expression and piRNA repertoires in relatively closely-related species with a focus on the 13-lined ground squirrel that appears to lack mobilizing LINE and SINE families. We found that squirrel TEs were still highly expressed but lacked a strong PIWI response.

### PIWI proteins are expressed later in squirrels

Mouse PIWIs, specifically PIWL2 and PIWIL4, are initially expressed in prenatal testis, however PIWIL4 ceases to be transcribed ~three days after birth but PIWIL2 is continuously expressed. Mice develop a little faster than squirrels, i.e. three vs four week pregnancy period, eyes open at P11-12 vs P21-24, and male mouse puberty occurs at four weeks while squirrels go through a hibernation cycle and do not reach puberty until 11 months after birth. Therefore, we wanted to address when PIWIs first become expressed in squirrel. We analyzed testis samples from squirrels between P0 and P13, and found that PIWIs are not expressed at least until P8. We acknowledge a six-day gap between P2 and P8 when PIWIs could have become expressed, and we lack information on embryonic expression. Consistently, the hallmarks of piRNA biogenesis and TE processing were not observed until P8 in squirrel, but unlike adult rabbit and mouse, the distribution of piRNAs in adult squirrel testis stayed centered around 28 nts instead of 30 nts, confounding the difference between prepachytene and pachytene piRNAs in this species. It is likely that the squirrel’s prolonged retention of PIWIL4 through adulthood contributed a higher abundance of short piRNAs.

Since PIWIs are not expressed in P0 and P2, the small RNAs examined cannot be piRNAs, but once PIWIs were expressed at P8, TEs became a more common source of these small RNAs (Fig 2B), many of which would be piRNAs. There were dramatic shifts in the sources of piRNAs between juvenile and adults of rabbits and mice, exhibiting differences in the sources of prepachytene and pachytene piRNAs, but this shift was not as apparent in squirrel. There could be two possible explanations for this: 1) because PIWIL4 is expressed in adult testis, TEs remain a common source of piRNAs or 2) TE transcripts are not heavily targeted by PIWI proteins and do not make up a considerable proportion of prepachytene piRNAs. Perhaps some combination of both is also possible. Consistently among the species analyzed, when PIWIL1 became expressed, diverse piRNAs synthesized from intergenic space became more abundant (Supplemental Table 1; Fig 2B).

### PIWI proteins did not reduce TE transcripts in squirrels

We initially predicted that TE expression variation among species could be attributed to the number of insertions in the genome. While this appeared to be true within genomes (Supplemental Figure 5), this was not the case among genomes (Figure 2A). Given the small percentage of the squirrel genome derived from TEs, especially young TEs, we expected TEs would make up a small proportion of the squirrel transcriptome, but in fact TEs made up the highest percentage of the transcriptome among species. This example of a disconnect between TE insertions and their expression is consistent with the idea that transcription and successful reintegration are distinct phenomena; the first does not necessarily lead to the second, and insertion success may not be dependent on high levels of TE expression (Deininger et al. 2003; Kazazian 2004; Lu and Clark 2010). However, given the apparent absence of retrotransposition-competent L1 loci in the squirrel assembly (Platt and Ray 2012), the lack of new integrations while still observing transcription was unsurprising.

There was some TE expression variation among the squirrel life stages revealed by PCA, where samples fell into three groups: a pre-PIWI group (P0 and P2), a post-PIWI juvenile group (P8, P10, and P13), and the adult group. However, TE expression changes could not be convincingly explained by the activity of PIWI proteins. Expression did not significantly change for 97.8% of TE families after PIWIs became expressed. Further, among the 17 families that were differentially expressed, seven were expressed higher in samples with expressed PIWIs. We yielded a similar result when we restricted analyses to young insertions, where 94.7% of families were not differentially expressed. However, of the eight differentially expressed families, seven experienced a reduction in samples with expressed PIWIs. Almost all the differentially expressed families were LTRs, and LTRs were the only group of elements with a strong ping-pong signature. Unfortunately, we could not perform the same test in mouse and rabbit for comparison, but it has been reported that L1s are 5-10x more expressed in mouse PIWIL2 knockouts (Aravin, Sachidanandam, et al. 2007). The absence of differentially expressed TEs could be explained by a lack of PIWI targeting, particularly in LINEs and SINEs. The strength of the ping-pong cycle was significantly reduced among squirrel LINEs and SINEs relative to rabbit and mouse (Fig 4A; B). We should note that the two most recently active L1s in squirrel had Z_10_ scores between 2.3 and 8.5, indicating an active ping-pong cycle, but substantially less so than the highest LINE ping-pong signatures in mouse and rabbit (max Z_10_ = 33.6 mouse, max Z_10_ = 81.3 rabbit). In a final approach to measure any LINE expression changes between squirrel juvenile and adult life stages, we examined ping-pong and LINE expression fold changes and obtained the same result. Neither expression nor ping-pong intensity changed substantially in the most recent squirrel LINE families (Fig 5).

A goal of ours was to test if PIWIs were responsible for the “death” of the squirrel L1. If that were the case, we predicted that the hallmarks of PIWI suppression as evidenced by an intense ping-pong cycle and low expression of targeted TEs, would still be present. However, we observed the opposite relationship. If PIWIs were responsible for the immobility of LINEs, those hallmarks are gone and PIWIs respond to these L1s like an old, harmless family, so the question remains: how do TE’s “die” and can PIWIs be responsible? Another unanswered question that remains is why TE expression is still so high in squirrels. One possible explanation could be due to the function of PIWI proteins. If there has been any functional relaxation in PIWIs, we should observe such a relaxation of functional constraint in the DNA sequences. To rule out this possibility, we collected PIWI paralog sequences from a range of Glires species and measured *d*_N_/*d*_S_ changes along the squirrel lineage. We found no evidence of a relaxation effect, but indeed found an intensification of purifying selection on PIWIL1. Further, there were still relatively strong ping-pong signatures among LTR families, indicating appropriate function. From these observations, we can rule out an explanation where PIWI proteins are at fault. Alternatively, Pasquesi et al (2020) demonstrated that the majority of vertebrate TE transcripts are derived from older TE insertions and Molaro et al. (2014) determined that mouse L1 promoters least diverged from consensus were methylated more heavily than promoters from older L1s. Further, Byun et al. (2013) found older TEs in the human genome are less methylated at CpG cites than younger families. Therefore, since the squirrel genome is largely lacking young TEs, it is plausible these TEs are not as methylated and transcribed at higher levels. However, this does not necessarily explain why young squirrel L1 insertions lack a strong ping-pong signature. This leads to the last major question that has yet to be answered is: do PIWIs distinguish between an expressed TE and a mobilizing TE and, if so, how? The squirrel and rabbit genomes contain a similar number of young L1s that are nearly equivalently expressed, but demonstrate a difference in the strength of the ping-pong cycle (Fig 4C). PIWIs are capable of targeting and cleaving TE transcripts from young families in a cluster-independent mechanism (Aravin et al. 2008; Gainetdinov et al. 2017), but these models do not explain how PIWIs are initially guided to the most threatening families, or how PIWIs differentiate between threatening and non-threatening transcripts.

Taken together, our results raise interesting questions regarding the interplay between TEs, piRNAs, and PIWI paralogs. Contrary to our expectations, we found that TEs were actively transcribed in the squirrel even though most are not accumulating in the genome. Squirrel PIWI proteins largely did not have an effect on the expression of most TEs and, unlike recently active LINEs in the two other mammals we examined, the ping-pong cycle did not lead to the reduction of TE transcripts. We hypothesize that this derives from the passive processing of TE transcripts into piRNAs in the squirrel, which sharply contrasts with the PIWI/piRNA associated active reduction of TE transcripts in the rabbit, mouse, and other mammals screened to date.

## Materials and Methods

### Sample acquisition and RNA sequencing

All animal procedures conformed with federal and institutional guidelines for humane care and use and were pre-approved by the respective Institutional Animal Care and Use Committees. 13-lined ground squirrels were reared in the captive breeding colony at the University of Wisconsin, Oshkosh (Merriman et al. 2012). For testis collection, euthanasia by cervical transection was conducted on neonates (P0), post-natal juveniles (P2, P8, P10, P13), and adults. We sampled one juvenile (P21) and one adult (P > 90) male rabbit. For each specimen, a cross section of testis was snap frozen in liquid nitrogen immediately following castration and stored at −80°C prior to RNA isolation. We isolated total RNA using Trizol ® (Invitrogen, USA) according to the manufacturer’s specifications. Small RNA libraries were prepared using the Illumina TruSeq small RNA kit © and 1 × 50 bp reads were sequenced on the Illumina Hi-Seq2000 platform. Directional RNASeq libraries were prepped using the Illumina TruSeq v2 kit and 2×100 bp reads were also sequenced on a HiSeq2000. All reads are available under the BioProject PRJNA528042. P7 mouse mRNA (SRR3659160), small RNAs (SRR3659150) adult mRNA (SRR765631) and small RNA (SRR772033) libraries were collected from the NCBI short-read archive (SRA).

### TE annotation

To visualize TE differences between the squirrel, rabbit and mouse genomes, we masked the ground squirrel (SpeTri2.0), rabbit (OryCun2.0.73) and mouse (GRCm38) genome using RepeatMasker 4.0.5 ‘-species *Ictidomys*’, ‘-species *Oryctolagus*’, and ‘-species *Mus*’, respectively. To estimate genetic distances, we used a modified calcDivergenceFromAlign.pl script included with RepeatMasker to calculate Kimura two-parameter (Kimura 1980) distances between each insertion and its respective consensus sequence. The option –noCpG was invoked to exclude highly mutable CpG sites from distance calculations. Redundant TE annotations were first identified using clusterBed (Quinlan and Hall 2010) and the longest annotation with the lowest K2P was selected as the most probable annotation for that locus (script available at https://github.com/mike2vandy/squirrel_piRNA).

### piRNA genomic mapping

Prior to small RNA mapping, we clipped barcodes, removed reads that had bases with Phred quality score <25, and removed identical reads using modules in the fastx toolkit. We also removed low complexity small RNA sequences by zipping the sequence and removing sequences that compressed by more than 75% (script available at https://github.com/mike2vandy/squirrel_piRNA). We mapped piRNA sequences 24-32 nts long to the respective genomes using Bowtie 1.2 (Langmead et al. 2009) with the parameters –a –v 0 to allow reads to map to all perfect matches in the genome. We estimated the source (LINE, SINE, LTR, DNA, RNA, genic, or other) of piRNAs given the most frequently mapped annotation type per piRNA. Further, we examined the strength of the ping-pong cycle among TE families. We used intersectBed to intersect piRNAs with TE annotations. For each TE family, we calculated a 10 nt complementary overlap Z_10_score where 1-9 and 11-20 nt overlaps were used as background following Han. et al (2015). We recorded the number of mapped positions for each small RNA and normalized piRNA abundances to all genome mapping reads as parts per million (ppm) so that our Z_10_score takes into account piRNA source ambiguity and piRNA amplification. We repeated the same analysis for insertions that were less than 5% diverged from TE consensus sequences. Scripts available at https://github.com/mike2vandy/squirrel_piRNA.

### TE expression

Prior to mapping, we removed adapters and poor-quality sequences from RNAseq data files using Trimmomatic 0.36 (Bolger et al. 2014). To estimate TE expression profiles through testis development, we mapped clean RNASeq reads to the appropriate genome draft and Ensembl annotated gtf using STAR v2.7.1a (Dobin et al. 2013) with the parameters --outFilterMultimapNmax 100 and --winAnchorMultimapNmax 100 to allow reads to map up to 100 positions. We then used TEtranscripts 2.0.3 (Jin et al. 2015), specifically TEcount, to simultaneously estimate gene and genome wide TE family expression from genome mapped RNAseq reads. TEcount requires two annotation files that specify gen and repeat coordinates. We used Ensembl annotations for genic coordinates and we modified our RepeatMasker output for compatibility with TEcount. Raw read counts were transformed into RPM counts using DESeq2 v1.26 (Love et al. 2014). We conducted a separate TE expression analysis to examine the expression patterns of just the “young” insertions. To do so, we filtered the TE gtf file for insertions that exhibited a Kimura 2-parameter distance less than 5% diverged from the consensus and reran TEcount.

To compare expression profiles among squirrel samples, we normalized read counts given all genes and TEs using estimateSizeFactors and estimateDispersions functions in DESeq2 after removing genes and TE families with less than 10 mapped reads across all samples. Patterns of expression variation were assessed by performing PCAs based on the blind variance stabilizing transformed data. We conducted PCAs with all genes and TEs, just TEs, and young TE insertions. We also performed a differential expression analysis between samples from juveniles that lacked expressed PIWI proteins (P0 and P2) and juveniles that expressed PIWI proteins (P8, P10, and P13) to identify TEs that were differentially expressed in the presence or absence of PIWI proteins. In these analyses, all TE families and genes were tested for differential expression and we determined statistical significance when the adjusted p-value was < 0.1. We conducted these analyses for a dataset that contained TE expression estimated from all TE insertions and a dataset based on the expression of young TE insertions. We performed log fold shrinkage for lowly expressed TEs using apeglm when plotting fold change against expression (Zhu et al. 2019).

### Selection tests

For each PIWI paralog, we collected orthologs from closely related rodent and lagomorph species from Ensembl (species and gene IDs are available in Supplemental Table 2). A codon alignment was made by translating coding sequences to amino acid, aligning amino acids using LINSI parameters in MAFFT (Katoh and Standley 2013), and using a custom script to reverse translate resulting alignments. Unrooted maximum likelihood trees were constructed using RaxML (Stamatakis 2014). We used the RELAX module (Wertheim et al. 2015) in HyPhy (Pond et al. 2005) to test for a relaxation or intensification of selection of the PIWI orthologs in squirrel. RELAX identifies shifts in the stringency of natural selection by performing a likelihood ratio test (LRT) between a null model that constrains a relation parameter K to 1 for all branches and an alternative model where K is a free parameter. K > 1 implies an intensification of selection and K < 1 suggests relaxation of selection along the test branch(es).

## Acknowledgments

We acknowledge the support from the National Science Foundation (DEB-1355176, RoL 1838283). Additional support was provided by the College of Agriculture and Life Sciences at Mississippi State University and the College of Arts and Sciences at Texas Tech University. In addition, we would like to thank the Texas Tech High-Performance Computing Center and the Mississippi State University Institute for Genomics Biocomputing and Biotechnology for providing the computational resources. Lastly, this work could not have been completed without the helpful assistance of Ildar Gainetdinov.

