## Supplementary material for "The PIWI/piRNA response is relaxed in a rodent that lacks mobilizing transposable elements": supplementalFigs.docx

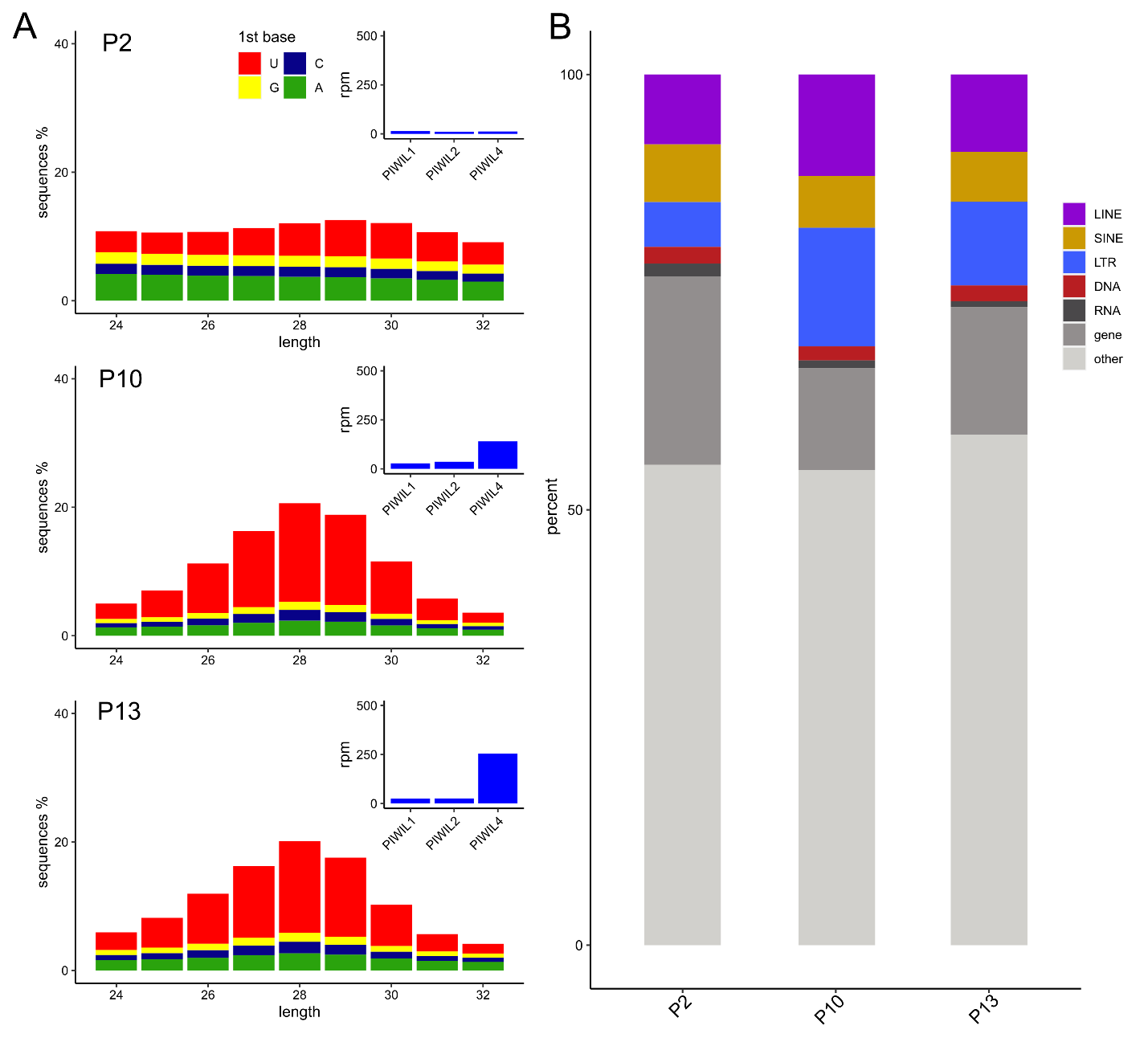

**Supplemental Figure 1**. A) PIWI expression, small RNA distributions and B) small RNA mapping frequencies of samples not included in Figure 2.

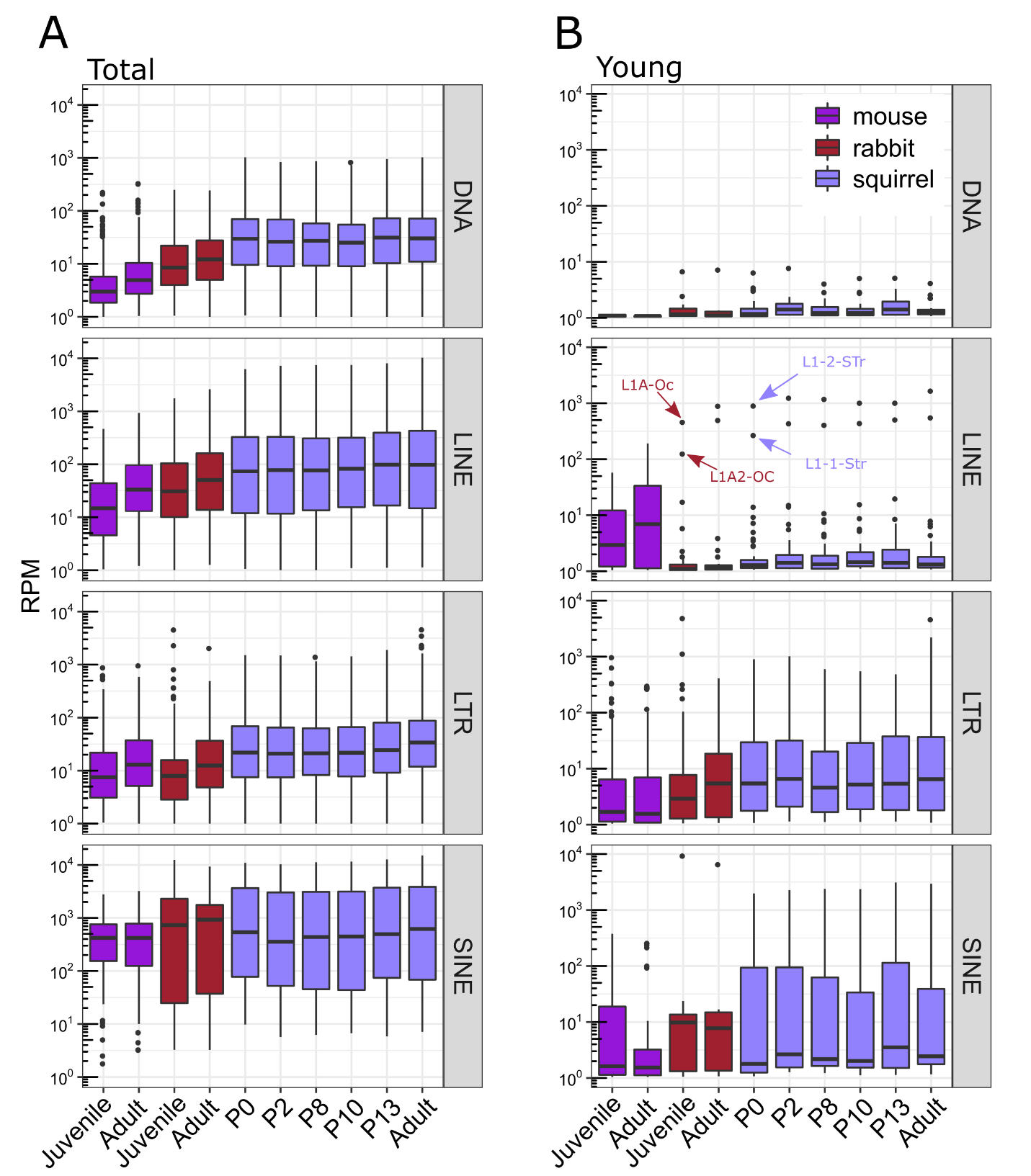

**Supplemental Fig 2**. A) TE expression distribution of families among species. B) TE family expression distribution calculated from young insertions.

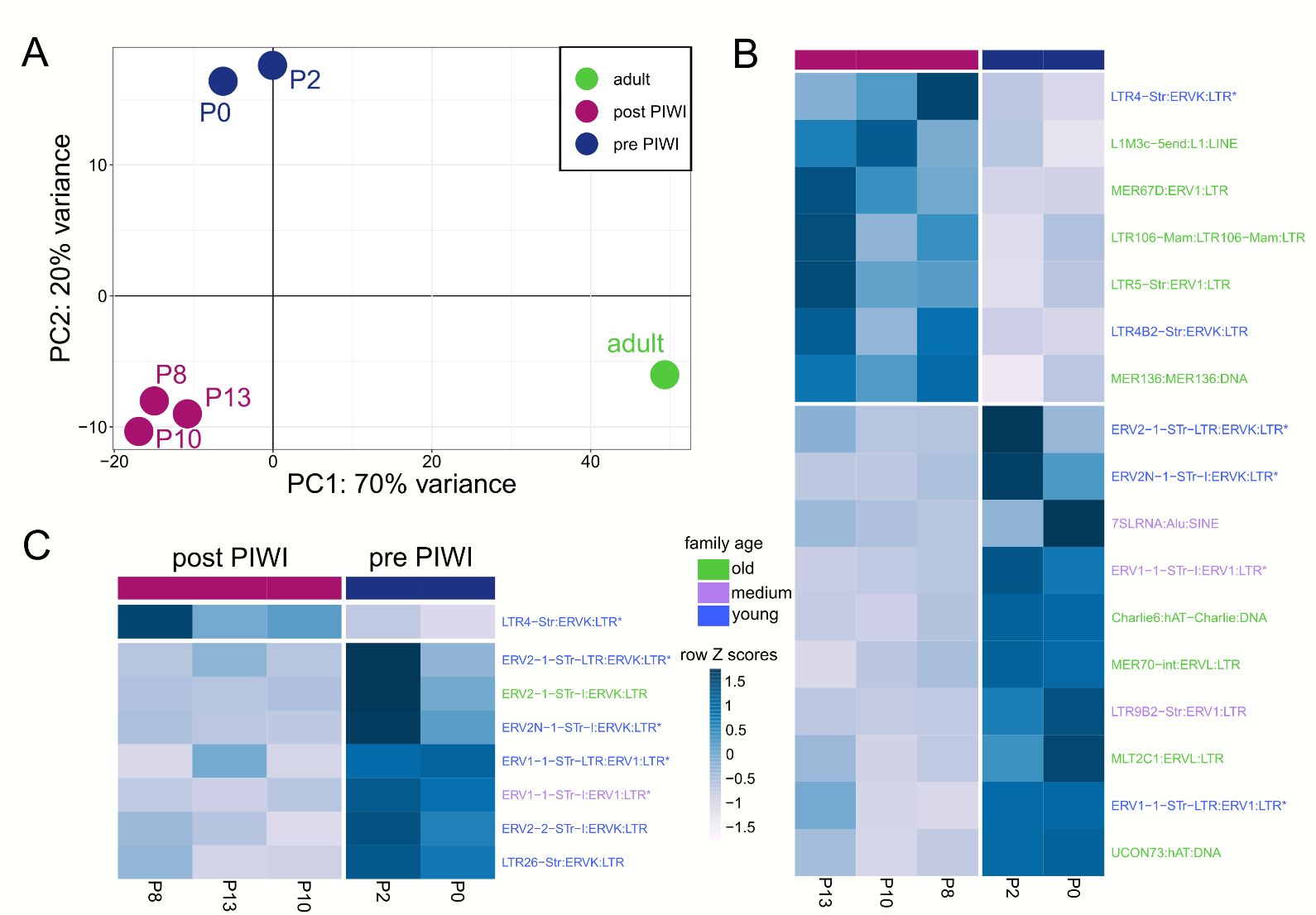

**Supplemental Fig 3**. A) PCA derived from the expression of all TEs and genes. Differentially expressed TEs between pre PIWI and post PIWI samples calculated from B) all TE insertions and C) young insertions. Specific families are color coded by age. Young families are <10% diverged from the consensus, medium are 10-20% diverged and old families are > 20% diverged.

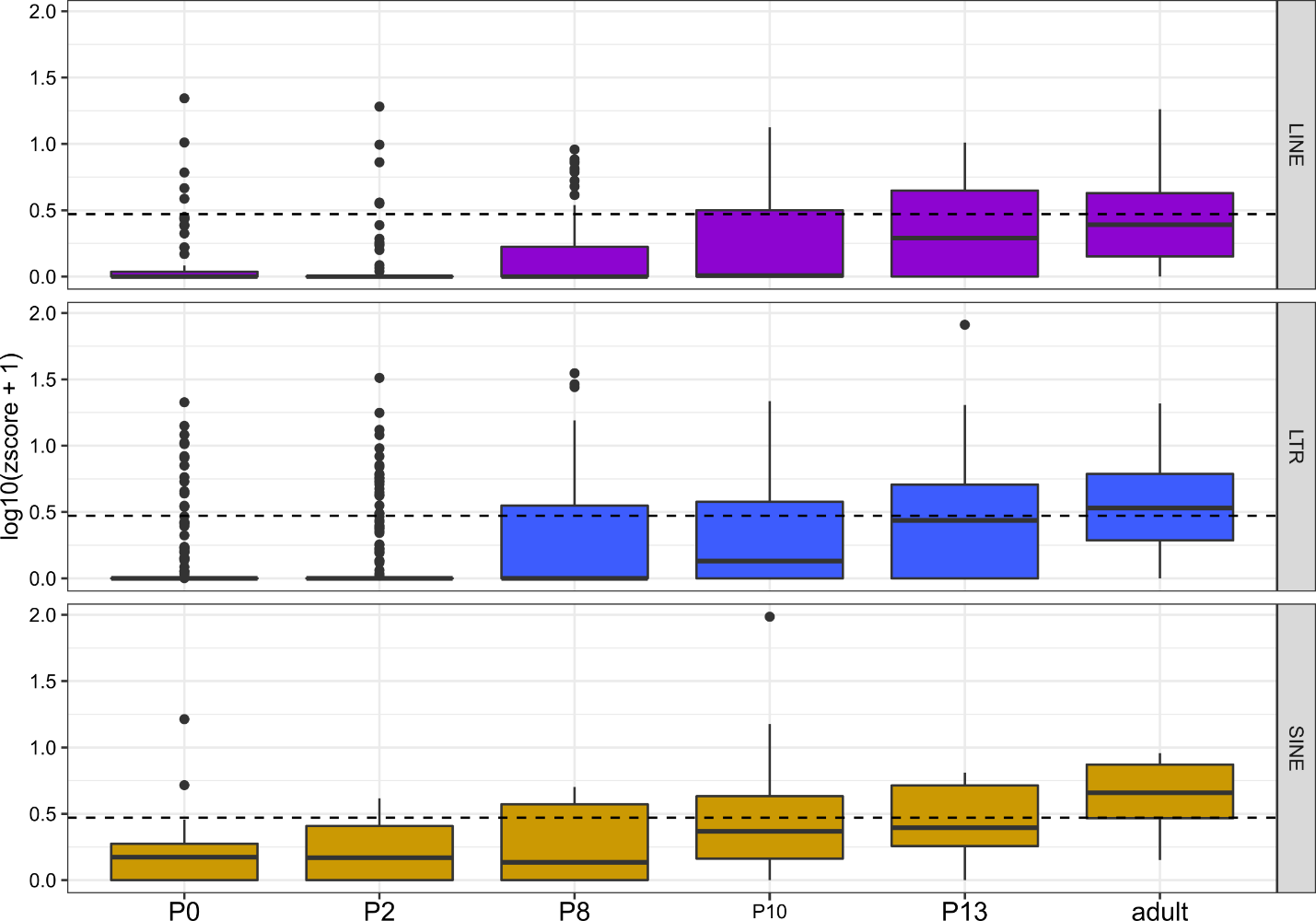

**Supplemental Fig 4**. Z10score distribution from TE families among the six life stages of squirrel. The dotted line reflects a Z_10_score = 1.95 p-value = 0.05.

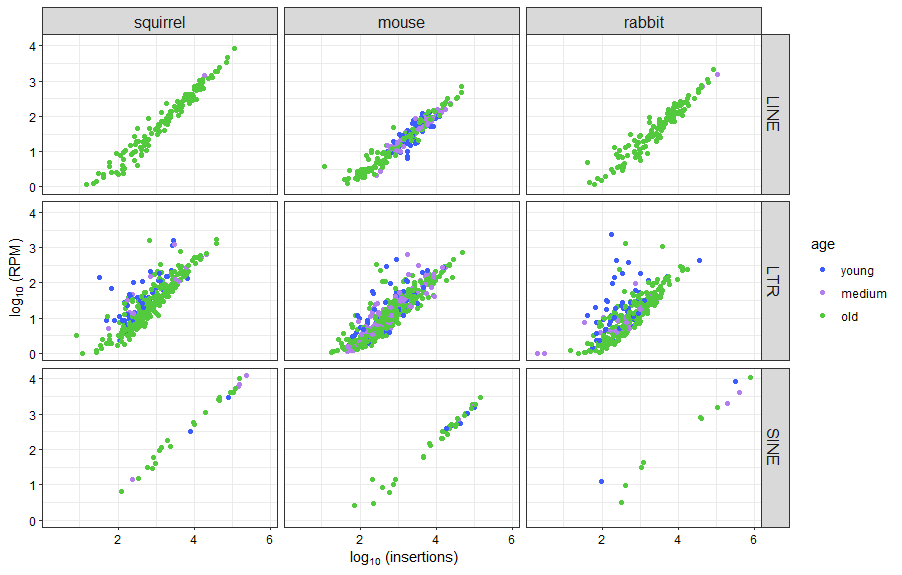

**Supplemental Fig 5**. Number of insertions per family plotted against family expression. Families are color coded by age where young families are <10% diverged from the consensus, medium are 10-20% diverged and old families are > 20% diverged.

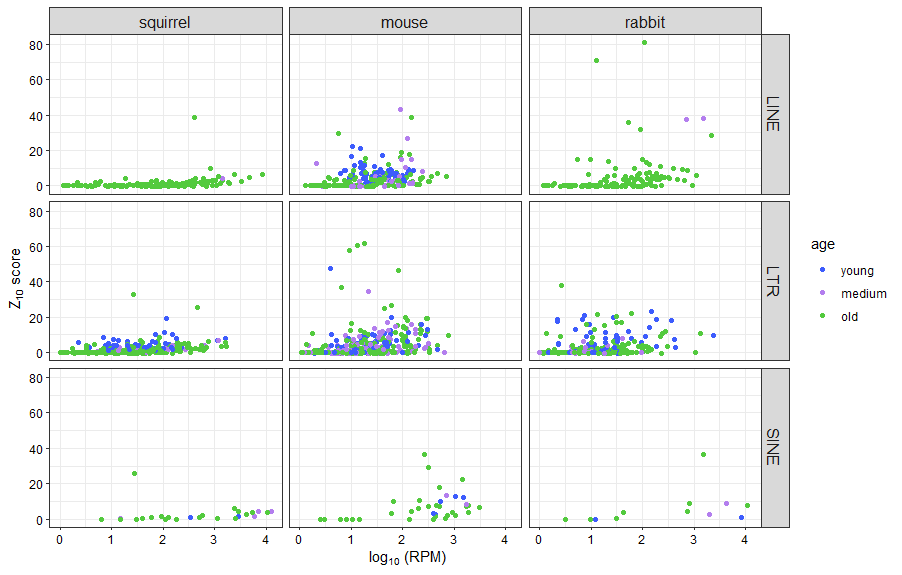

**Supplemental Fig 6.** TE family expression plotted against TE family Z_10_scores. Families are color coded by age where young families are <10% diverged from the consensus, medium are 10-20% diverged and old families are > 20% diverged.

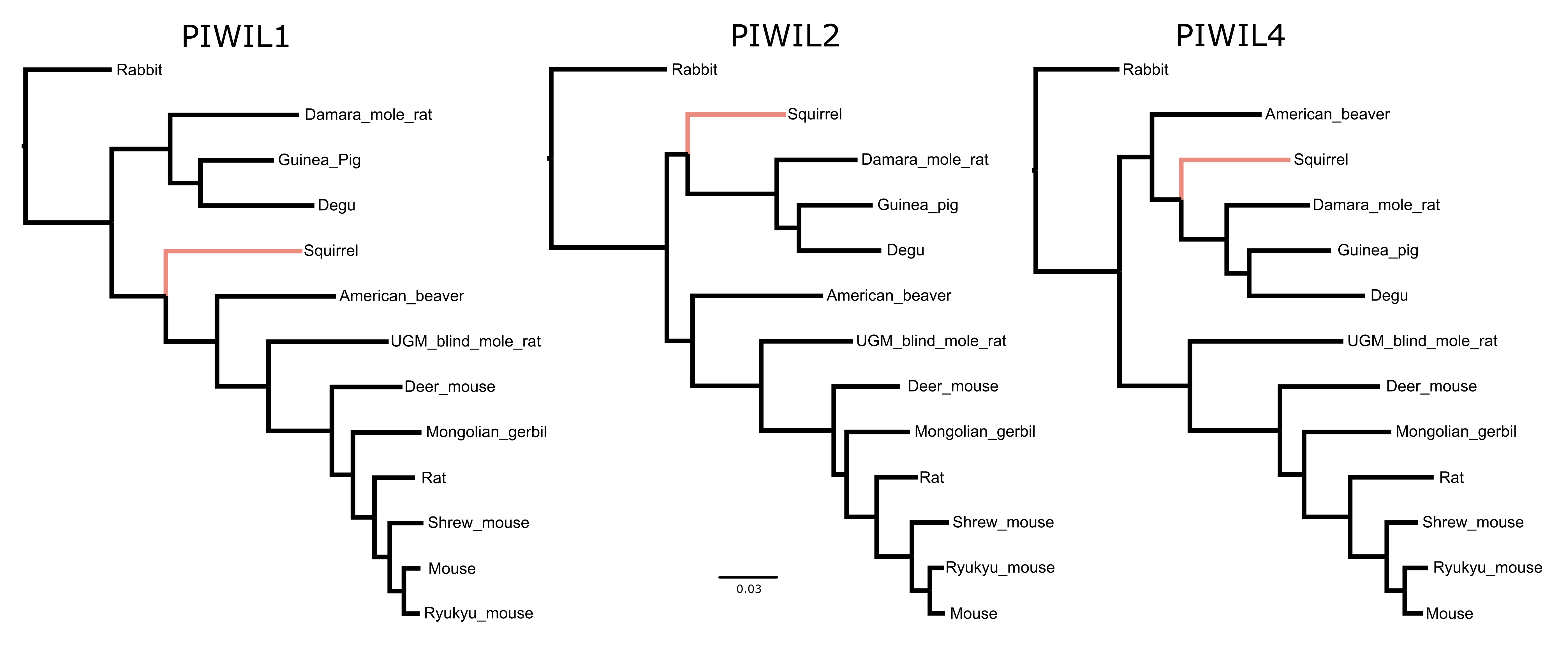

**Supplemental Fig 7**. PIWI gene trees used for RELAX tests in HyPhy. Trees presented here are rooted for easier viewing but were not rooted in selection tests. The pink branch was the labeled test branch.
